## Supplemental Figures, Figure Legends and Movie Legends for "ICAM-1 nanoclusters regulate hepatic epithelial cell polarity by leukocyte adhesion-independent control of apical actomyosin"

**Cacho Navas et al.**

### SUPPLEMENTARY FIGURES, MOVIES AND LEGENDS

#### SUPPLEMENTARY FIGURES AND FIGURE LEGENDS

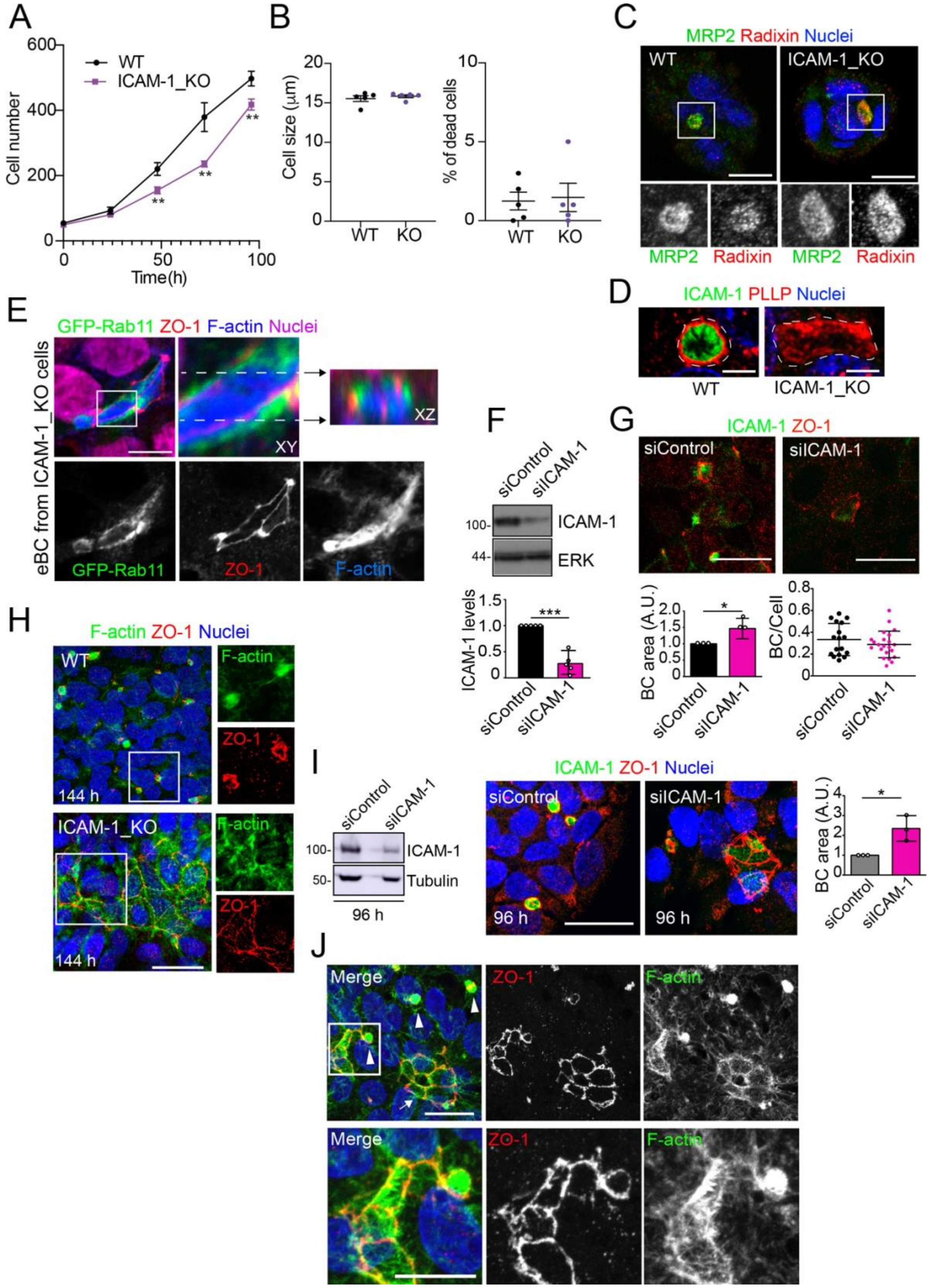

**Supplementary Figure 1. Related to Figures 1 and 2. (A)** Cells were seeded at very low density and counted for 96 h. Bars represent the SD of the mean. n=3. ** p<0.01. **(B)** In each cell passage, cell density, size (left graph) and percentage of dead cells (right graph) were analyzed in a cell counter Countess 3 (Invitrogen). Bars represent the SD of the mean. n=5. **(C)** Control WT and ICAM-1_KO HepG2 cells were cultured on coverslips for 72 h, fixed with methanol at -20$^{o}$C for 5 min , and stained for MRP2 and radixin. Bottom images are single channel images of the squared areas. Scale bars, 10 µm. **(D)** Control WT and ICAM-1_KO HepG2 cells were cultured on coverslips for 72 h, fixed with formalin 10% at room temperature for 20 min and stained for plasmolipin/PLLP and F-actin. Scale bars, 5 µm. **(E)** Image of ICAM-1_KO HepG2 cells stably expressing GFP-Rab11 and stained for the indicated proteins (top left panel), in which the boxed area has been three-fold enlarged and its XY and XZ projections displayed in the top center and top right panels, respectively. Bottom panels show the top left image splitted into its three channels. Scale bar, 10 µm. **(F,G)** HepG2 cells were transfected with siRNA control (siControl) or with siRNA targeting ICAM-1 (siICAM-1). Cells were lysed and analysed by western blot with the indicated antibodies (F) or fixed, immunostained with the indicated antibodies and analyzed by confocal microscopy (G) Scale bar, 20 µm. (F) Quantification of ICAM-1 expression levels. A.U., arbitrary units. n=5. (G) Quantification of BC area (left graph) and frequency (BC/cell, right graph). n=3. **(H)** WT and ICAM-1_KO HepG2 cells were culture for 6 days, fixed and stained for ZO-1 and F-actin. Right images show enlargements of the boxed areas in the left images. Scale bar, 20 µm. **(I)** HepG2 cells were transfected with siRNA control or with siRNA targeting ICAM-1 and cultured for 96 h. Cells were lysed and subjected to western blot analysis with the indicated antibodies (left panels) or fixed and analyzed by immunofluorescence and confocal microscopy (central images). Tubulin is shown as a loading control. The right graph shows the quantification of BC area. Scale bar, 20 μm. Bars represent the mean ± SD. n=3. * p<0.05; *** p<0.001. A.U. Arbitrary Units. Nuclei were stained with DAPI. **(J)** Distribution of F-actin and ZO-1 in ICAM-1_KO cells cultured for 96 h suggest coalescence of small BCs into eBC. Top images. Arrowheads point to BCs and the arrow to an eBC. Bottom images corresponded to two-fold enlargement of the top squared area and show a F-actin-enriched spherical BC in contact with a massive eBCs. Scale bars. Top, 20 μm; bottom, 10 μm. Bars represent the mean ± SD. n≥ 3. * p<0.05; *** p<0.001. A.U. Arbitrary Units. Nuclei were stained with DAPI.

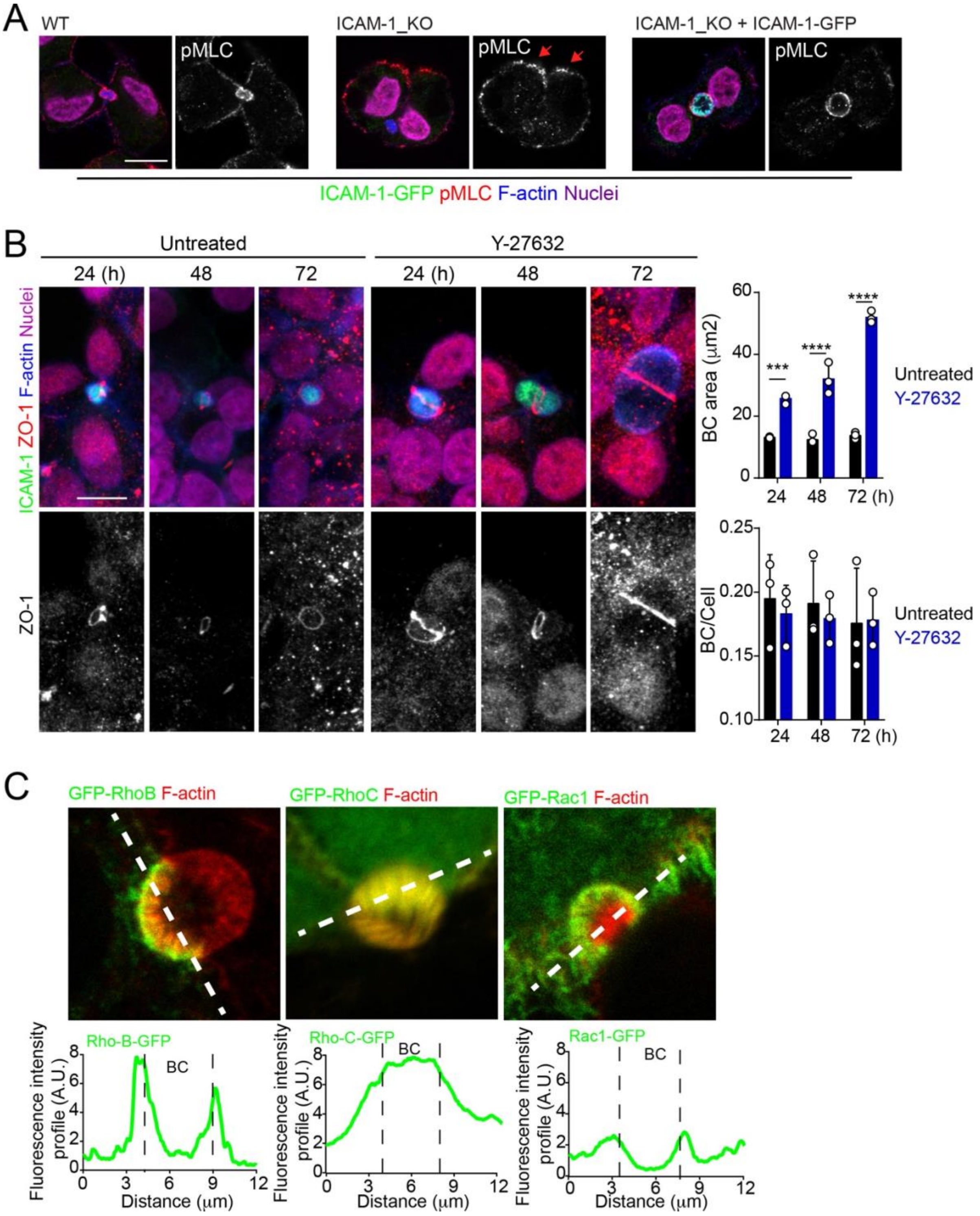


**Supplementary Figure 2. Related to Figure 4**. **(A)** Representative images of cells quantified in Figure 4D. **(B)** HepG2 cells were exposed or not to 5 μM of the ROCK inhibitor Y27632 for the indicated times. Scale bar, 10 μm. Graphs shows the quantification of the effect of

Y27632 on BC area (top) and BC frequency (BC/cell, bottom). At least 20 BCs were quantified in each of the three experiment. **(C)** Cells were transfected and the expression of the indicated GFP-Rho proteins analyzed 24 h post-transfection. The GFP intensity profiles along the dotted lines are shown in the bottom graphs.

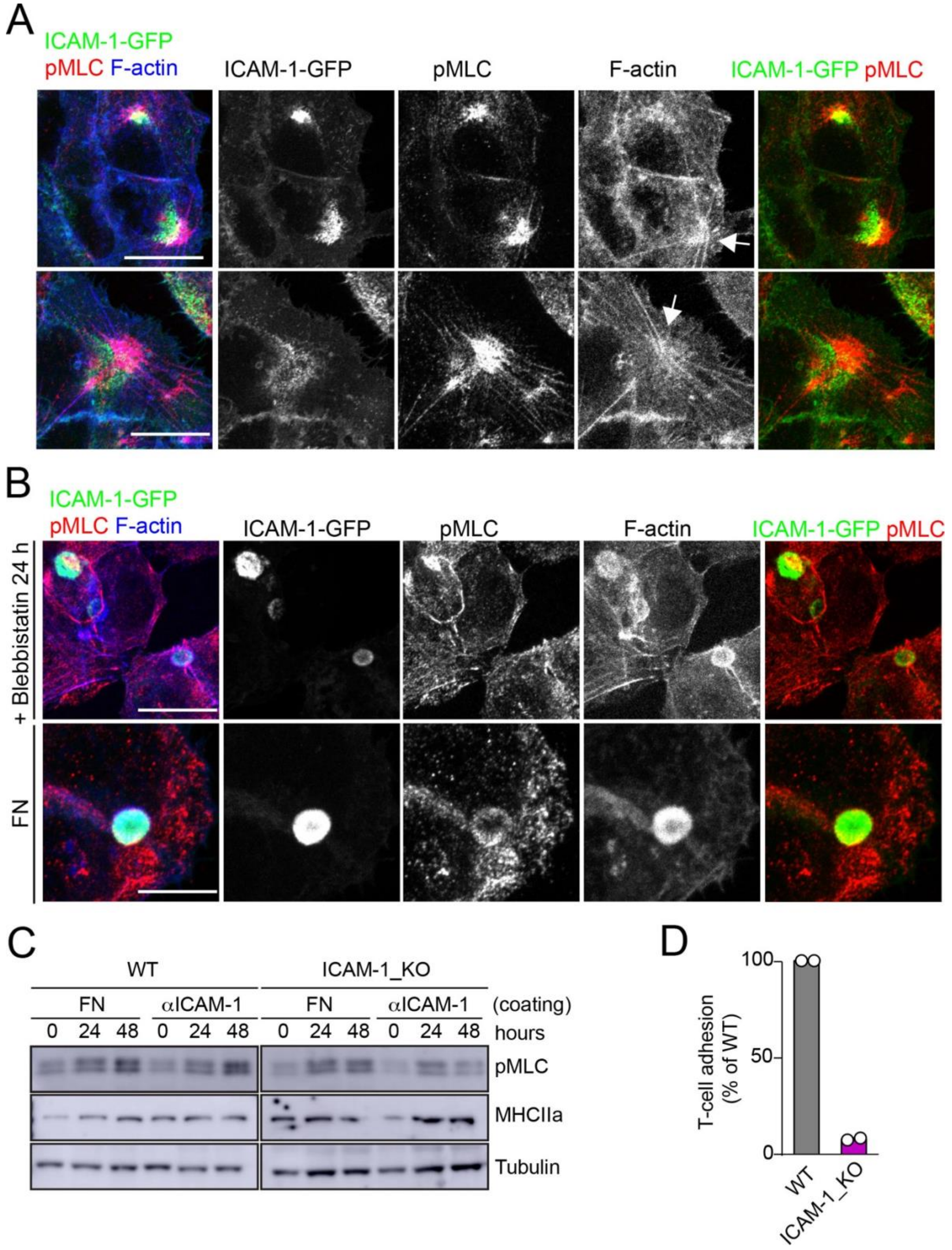


**Supplementary Figure 3. Related to Figure 5. (A)** Surface ICAM-1 engagement generates stellate stress fibers that concentrate pMLC and ICAM-1, although in different cellular domains. ICAM-1-GFP HepG2 cells were cultured for 24 h on coverslips precoated with anti-

ICAM-1. Cells were fixed and stained for the indicated proteins. Two representative images of stellate stress fibers (arrows) and their corresponding splitted channels are shown. **(B)** Cells on coverlsips coated with anti-ICAM-1 were incubated with 10 μm blebbistatin before fixation (top). In parallel, cells were plated on FN-coated converslips for 24 h (bottom). Cells were fixed and stained as in (A) Scale bars, 10 μm. **(C)** WT and ICAM_KO cells were seeded on plates precoated with FN or anti (α)-ICAM-1 antibody for the indicated times, lysed and subjected to western blot to detect the indicated proteins. **(D)** CD3-positive T-cells were quantified in experiments performed as in Figure 5F. WT and ICAM-1_KO cells were cultured for 72 h and incubated with T-lymphocytes for 6 h. The graph shows the percentage of T-cell adhesion with respect to WT cells in two different experiments.

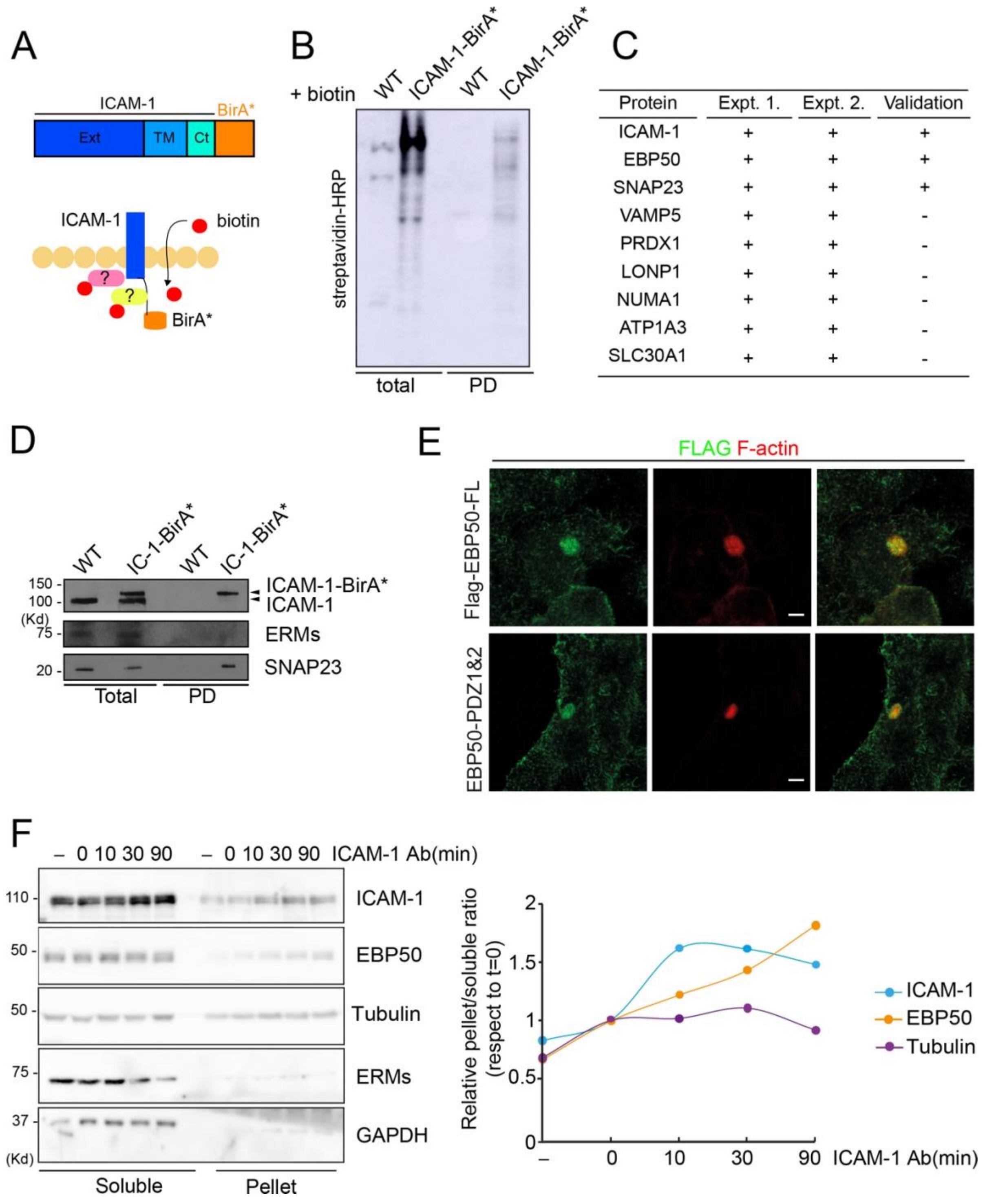


| Protein | Expt. 1. | Expt. 2. | Validation |
|---|---|---|---|
| ICAM-1 | + | + | + |
| EBP50 | + | + | + |
| SNAP23 | + | + | + |
| VAMP5 | + | + | - |
| PRDX1 | + | + | - |
| LONP1 | + | + | - |
| NUMA1 | + | + | - |
| ATP1A3 | + | + | - |
| SLC30A1 | + | + | - |

**Supplementary Figure 4. Related to Figure 6**. ICAM-1 BioID reveals the proximal interaction of the receptor with a new set of proteins. **(A)** Schematic representation of the proximal biotinylation of proteins by the ICAM-1-BirA* chimeric protein. **(B)** Cells expressing ICAM-1-

BirA* or not were incubated with 50 μM biotin for 16 h, lysed and subjected to a pull-down (PD) assay with neutravidin-agarose. Biotinylated proteins were detected by blot with streptavidin-HRP. **(C)** Pulldown fractions from two experiments were subjected to mass spectrometry analysis. Table shows the proteins identified exclusively in the fraction of HepG2 expressing ICAM-1-BirA* but not in the pulldown fraction of parental cells. EBP50 proximal interaction with ICAM-1 is validated by western blot in Figure 6a **(D)** Proximal interaction of SNAP23, but not of ezrin-radixin-moesin proteins, despite both have been previously described as ICAM-1 interactors (5,42). **(E)** Cells stably expressing the indicated Flag-tagged EBP50 proteins were cultured for 48h, fixed and stained for the FLAG epitope and for F-actin to localized BCs. Scale bars, 5 μm. **(F)** Cells were incubated with anti-ICAM-1 mAb for 30 min at 4$^{o}$C, washed and incubated at 37$^{o}$C for the indicated times. Cells were lysed and soluble and insoluble (pellet) fractions were separated by centrifugation as described in Materials and Methods. The relative distribution of the indicated proteins was analyzed by western blot (left) and quantified (right).

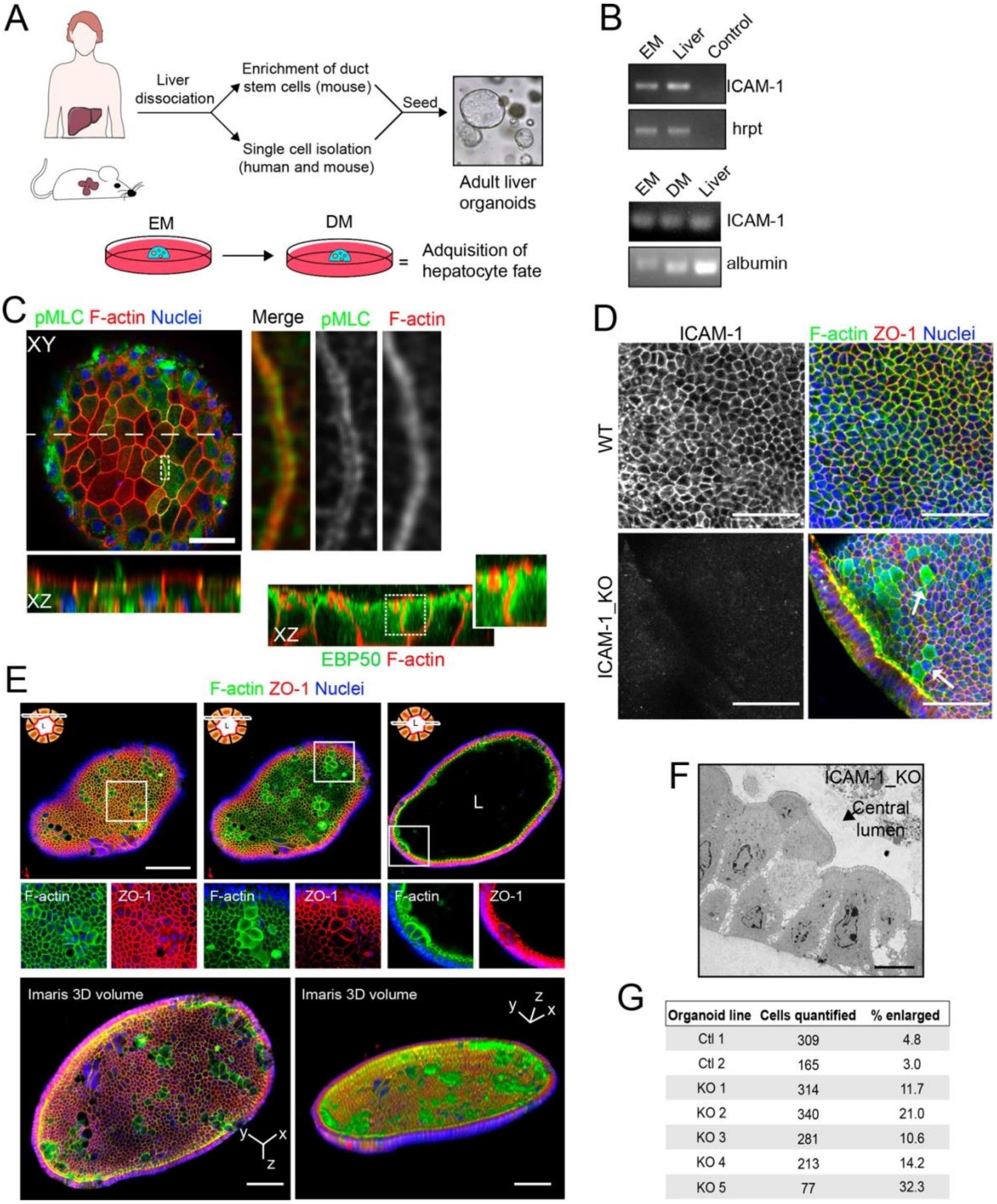


| Organoid line | Cells quantified | % enlarged |
|---|---|---|
| Ctl 1 | 309 | 4.8 |
| Ctl 2 | 165 | 3.0 |
| KO 1 | 314 | 11.7 |
| KO 2 | 340 | 21.0 |
| KO 3 | 281 | 10.6 |
| KO 4 | 213 | 14.2 |
| KO 5 | 77 | 32.3 |

**Supplementary Figure 5. Related to Figure 8**. **(A)** Graphical protocol of ductal stem cell isolation and differentiation from human and murine liver biopsies (Broutier et al. 2016). Briefly, ductal stem cells were isolated from hepatic tissue and place in a 3D environment in the presence of expansion medium (EM). Differentiation to the hepatocyte lineage was induced by changing EM to differentiation medium (DM) for 13 days. **(B)** PCR analysis of ICAM-1 and albumin mRNA in murine liver organoids and liver tissue. A PCR control with no primers is shown. **(C)** Murine liver organoids were fixed, permeabilized and stained for pMLC, F-actin and nuclei. Left image show a confocal plane that crosses the central part of an epithelial sheet of the organoid. Bottom image shows the XZ reconstruction of this image in the plane labeled with a discontinuous line. Right images show an eight-fold magnification of the boxed area showing that pMLC surrounds lateral, perijunctional F-actin domains. Bottom right images show the XZ reconstruction of a murine liver organoid stained for F-actin and EBP50, in which EBP50 is enriched in perijunctional areas. **(D)** Z-stack projection of confocal images taken from WT and ICAM-1_KO organoids in which the size of some ICAM-1_KO cells is expanded (arrow). Scale bar, 50 μm. **(E)** Differentiated ICAM-1_KO organoids were fixed and stained for F-actin and ZO-1. Top images. Two different confocal planes in the Z-axis of the same organoid. Bottom images show a two-fold increase of the boxed area to show details of F-actin and ZO-1 distribution in the enlarged cells. Scale bars, 100 μm. Nuclei were stained with DAPI. **(F)** Transmission electron microscopy of ICAM-1_KO cells in which apical membranes protrude into the central lumen. Scale bar, 10 μm. **(G)** Quantification of EM images of WT and ICAM-1_KO organoids. Table shows the percentage of cells with enlarged morphology: abnormally enlarged and/or protruding from the cell monolayer that forms the organoid.

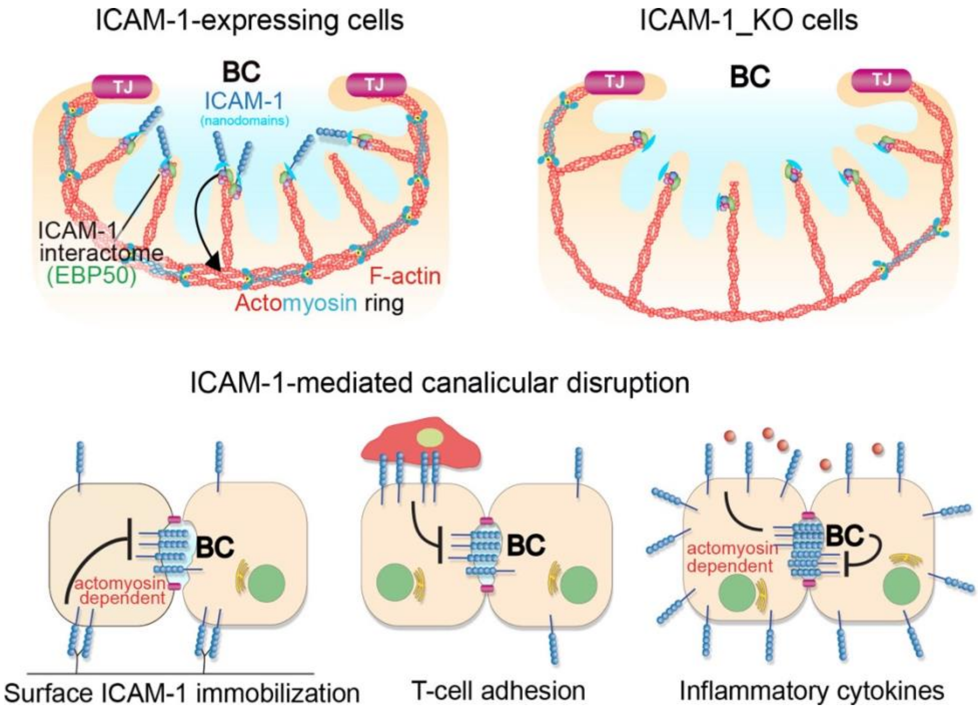


**Supplementary Figure 6.** A model for the control of actomyosin-mediated contraction of BCs mediated by ICAM-1 expression levels, which determine the size and frequency of these apical structures in hepatic epithelial cells. Top images. Canalicular ICAM-1 signals towards a distal canalicular actomyosin ring at the base of microvilli. EBP50 links ICAM-1 to F-actin and localizes into membrane nano-scale domains to mediate this signaling. In ICAM-1_KO actomyosin-mediated contraction is reduced and BCs are enlarged. Bottom images. ICAM-1 can signal towards actomyosin and regulate BC frequency. Surface immobilization of ICAM-1 prevents BC formation (bottom left) or disrupts mature BCs, as shown by ICAM-1-medated T-cell adhesion (bottom center). ICAM-1 upregulation in response to inflammatory cytokines increases actomyosin mediated contraction at BCs, which also reduces BC frequency (bottom right).

**SUPPLEMENTARY MOVIE LEGENDS**

**Movie 1. Related to Figure 2B.** Time-lapse fluorescence microscopy of polarized WT HepG2 cells stably expressing MDR1-GFP. Images were acquired at 15 min intervals for 18 h and displayed at 4 frame per second. Note the fusion of two BCs between 14 and 18 h. Scale bar 10 μm.

**Movie 2. Related to Figure 2B.** Time-lapse fluorescence microscopy of polarized ICAM-1_KO HepG2 cells stably expressing MDR1-GFP. Images were acquired at 15 min intervals for 18 h and displayed at 4 frame per second. Scale bar 10 μm.

**Movie 3. Related to Figure 4A.** Time-lapse spinning disc confocal microscopy of polarized HepG2 cells expressing GFP-tagged myosin light chain (GFP-MLC)(green) and incubated with SirActin (red) Images were acquired at 4 min intervals for 48 min and displayed at 1 frame per second. Scale bar, 3 μm.
